# pong: fast analysis and visualization of latent clusters in population genetic data

**DOI:** 10.1101/031815

**Authors:** Aaron A. Behr, Katherine Z. Liu, Gracie Liu-Fang, Priyanka Nakka, Sohini Ramachandran

**Affiliations:** Department of Ecology and Evolutionary Biology, Brown University, Providence, RI, USA; Department of Computer Science, Brown University, Providence, RI, USA; Computer Science Department, Wellesley College, Wellesley, MA, USA; Center for Computational Molecular Biology, Brown University, Providence, RI, USA

## Abstract

**Motivation:** A series of methods in population genetics use multilocus genotype data to assign individuals membership in latent clusters. These methods belong to a broad class of mixed-membership models, such as latent Dirichlet allocation used to analyze text corpora. Inference from mixed-membership models can produce different output matrices when repeatedly applied to the same inputs, and the number of latent clusters is a parameter that is often varied in the analysis pipeline. For these reasons, quantifying, visualizing, and annotating the output from mixed-membership models are bottlenecks for investigators across multiple disciplines from ecology to text data mining.

**Results:** We introduce `pong`, a network-graphical approach for analyzing and visualizing membership in latent clusters with a native D3.js interactive visualization. `pong` leverages efficient algorithms for solving the Assignment Problem to dramatically reduce runtime while increasing accuracy compared to other methods that process output from mixed-membership models. We apply `pong` to 225,705 unlinked genome-wide single-nucleotide variants from 2,426 unrelated individuals in the 1000 Genomes Project, and identify previously overlooked aspects of global human population structure. We show that `pong` outpaces current solutions by more than an order of magnitude in runtime while providing a customizable and interactive visualization of population structure that is more accurate than those produced by current tools.

**Availability:** `pong` is freely available and can be installed using the Python package management system `pip`. `pong`’s source code is available at https://github.com/abehr/pong.

## 5 Introduction

A series of generative models known as mixed-membership models have been developed that model grouped data, where each group is characterized by a mixture of latent components. One well-known example of a mixed-membership model is latent Dirichlet allocation (Blei <u>et al.</u>, 2003), in which documents are modeled as a mixture of latent topics. Another widely used example is the model implemented in the population-genetic program Structure (Pritchard <u>et al.</u>, 2000; Falush <u>et al.</u>, 2003; Hubisz <u>et al.</u>, 2009; Raj <u>et al.</u>, 2014), where individuals are assigned to a mixture of latent *clusters*, or populations, based on multilocus genotype data.

In this paper, we focus on the population-genetic application of mixed-membership models, and refer to this application as *clustering inference*; see Novembre (2014) for a review of multiple population-genetic clustering inference methods, including Structure. In Structure’s Bayesian Markov chain Monte Carlo (MCMC) algorithm, individuals are modeled as deriving ancestry from *K* clusters, where the value of *K* is user-specified. Each cluster is constrained to be in Hardy-Weinberg equilibrium, and clusters vary in their characteristic allele frequencies at each locus. Clustering inference using genetic data is a crucial step in many ecological and evolutionary studies. For example, identifying genetic subpopulations provides key insight into a sample’s ecology and evolution (Bryc <u>et al.</u>, 2010; Glover <u>et al.</u>, 2012; Moore <u>et al.</u>, 2014), reveals ethnic variation in disease phenotypes (Moreno-Estrada <u>et al.</u>, 2014), and reduces spurious correlations in genome-wide association studies (Price <u>et al.</u>, 2006; Patterson <u>et al.</u>, 2006; Galanter <u>et al.</u>, 2012).

For a given multilocus genotype dataset with *N* individuals and *K* clusters, the output of a single algorithmic run of clustering inference is an *N* × *K* matrix, denoted as *Q*, of *membership coefficients*; these coefficients can be learned using a supervised or unsupervised approach. Membership coefficient *q_ij_* is the inferred proportion of individual *i*’s alleles inherited from cluster *j*. The row vector 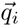. is interpreted as the genome-wide ancestry of individual *i*, and the *K* elements of x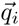. sum to 1. Each column vector 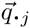 represents membership in the *j*th cluster across individuals.

Although covariates — such as population labels, geographic origin, language spoken, or method of subsistence — are not used to infer membership coefficients, these covariates are essential for interpreting *Q* matrices. Given that over 16,000 studies have cited Structure to date, and 100 or more *Q* matrices are routinely produced in a single study, investigators need efficient algorithms that enable accurate processing and interpretation of output from clustering inference.

Algorithms designed to process *Q* matrices face three challenges. First, a given run, which yields a single *Q* matrix, is equally likely to reach any of *K*! column-permutations of the same collection of estimated membership coefficients due to the stochastic nature of clustering inference. This is known as *label switching*: for a fixed value of *K* and identical genetic input, column 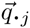 in the *Q* matrix produced by one run may not correspond to column 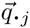 in the *Q* matrix produced by another run (Stephens, 2000; Jasra <u>et al.</u>, 2005; Jakobsson and Rosenberg, 2007). In our analyses of the 1000 Genomes (phase 3; Consortium (2015)), label switching occurred in 62.64 percent of pairwise comparisons among runs; that is, many matrices of membership coefficients were identical once columns were permuted to match, and rapidly finding permutations that maximize similarity between *Q* matrices is computationally expensive as *K* increases.

Second, even after adjusting for label switching, *Q* matrices with the same input genotype data and the same value of *K* may differ non-trivially. This is known as *multimodality* (Jakobsson and Rosenberg, 2007), and occurs when multiple sets of membership coefficients can be inferred from the data. We refer to runs that, despite identical inputs, differ non-trivially as belonging to different *modes.* For a fixed value of *K*, a set of runs grouped into the same mode based on some measure of similarity can be represented by a single *Q* matrix in that mode. Many studies using the maximum-likelihood approach for clustering inference implemented in ADMIXTURE (Alexander <u>et al.</u>, 2009) ignore manifestations of multimodality (Moreno-Estrada <u>et al.</u>, 2013; Consortium, 2015; Homburger <u>et al.</u>, 2015), despite the fact that ADMIXTURE can identify different local maxima across different runs for a given value of *K* (e.g., Verdu <u>et al.</u> (2014)). The complete characterization of modes present in clustering inference output gives unique insight into genetic differentiation within a sample.

A third complication arises for interpreting clustering inference output when the input parameter *K* is varied (all other inputs being equal): there is no column-permutation of a *Q*_*N* × *K*_ matrix that exactly corresponds to any *Q*_*N* × (*K* + * 1)_ matrix. We refer to this as the *alignment-across-K* problem. A common pipeline when applying clustering inference methods to genotype data is to increment *K* from 2 to some user-defined maximum value *K_max_*, although some clustering inference methods also assist with choosing the value of *K* that best explains the data (Huelsenbeck <u>et al.</u>, 2011; Raj <u>et al.</u>, 2014). *K_max_* can vary a great deal across studies (e.g., *K_max_* = 5 in Glover <u>et al.</u> (2012); *K_max_* = 20 in Moreno-Estrada <u>et al.</u> (2014)). Accurate and automated analysis of clustering inference output across values of *K* is essential both for understanding a sample’s evolutionary history and for model selection.

The label-switching, multimodality, and alignment-across-*K* challenges must all be resolved in order to fully and accurately characterize genetic differentiation and shared ancestry in a dataset of interest. Here, we present `pong`, a new algorithm for fast post-hoc analysis of clustering inference output from population genetic data combined with an interactive JavaScript data visualization using Data-Driven Documents (D3.js; https://github.com/mbostock/d3). Our package accounts for label switching, characterizes modes, and aligns *Q* matrices across values of *K* by constructing weighted bipartite graphs for each pair of *Q* matrices depicting similarity in membership coefficients between clusters. Our construction of these graphs draws on efficient algorithms for solving the combinatorial optimization problem known as the Assignment Problem, thereby allowing `pong` to process hundreds of *Q* matrices in seconds. `pong` displays a representative *Q* matrix for each mode for each value of *K*, and identifies differences among modes that are easily missed during visual inspection. We compare `pong` against current solutions (*CLUMPP* by Jakobsson and Rosenberg (2007); augmented as Clumpak by Kopelman <u>et al.</u> (2015)), and find our approach reduces runtime by more than an order of magnitude. We also apply `pong` to clustering inference output from the 1000 Genomes (phase 3) and present the most comprehensive depiction of global human population structure in this dataset to date. `pong` has the potential to be applied broadly to identify modes, align output, and visualize output from inference based on mixed-membership models.

## 6 Algorithm

### 6.1 Overview

Figure 1 displays a screenshot of `pong`’s visualization of population structure in the 1000 Genomes data (phase 3, Consortium (2015); final variant set released on November 6, 2014) based a set of 20 runs (*K* = 4, 5) from clustering inference with ADMIXTURE (Alexander <u>et al.</u>, 2009). In order to generate visualizations highlighting similarities and differences among *Q* matrices, `pong` generates weighted bipartite graphs connecting clusters between runs within and across values of *K* (Section 6.2). Our goal of matching clusters across runs is analogous to the combinatorial optimization problem known as the Assignment Problem (Manber, 1989), for which numerous efficient algorithms exist (Kuhn, 1955, 1956; Munkres, 1957). `pong`’s novel approach of comparing clusters — column vectors of *Q* matrices — dramatically reduces runtime relative to existing methods that rely on permuting entire matrices.

Consider two *Q* matrices, 𝓠 = [*q_ij_*] and 𝓡 = [*r_ij_*]. Each weighted bipartite graph G(𝓠, 𝓡) = 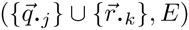 encodes pairwise similarities between clusters in 𝓠 and clusters in 𝓡. Edges in *G* are weighted according to a similarity metric computed between clusters (detailed in Supplementary Information); `pong`’s default similarity metric is derived from the Jaccard index used in set comparison, and emphasizes overlap in membership coefficients without incorporating individuals who have no membership in the clusters under comparison.

**Figure 1:**
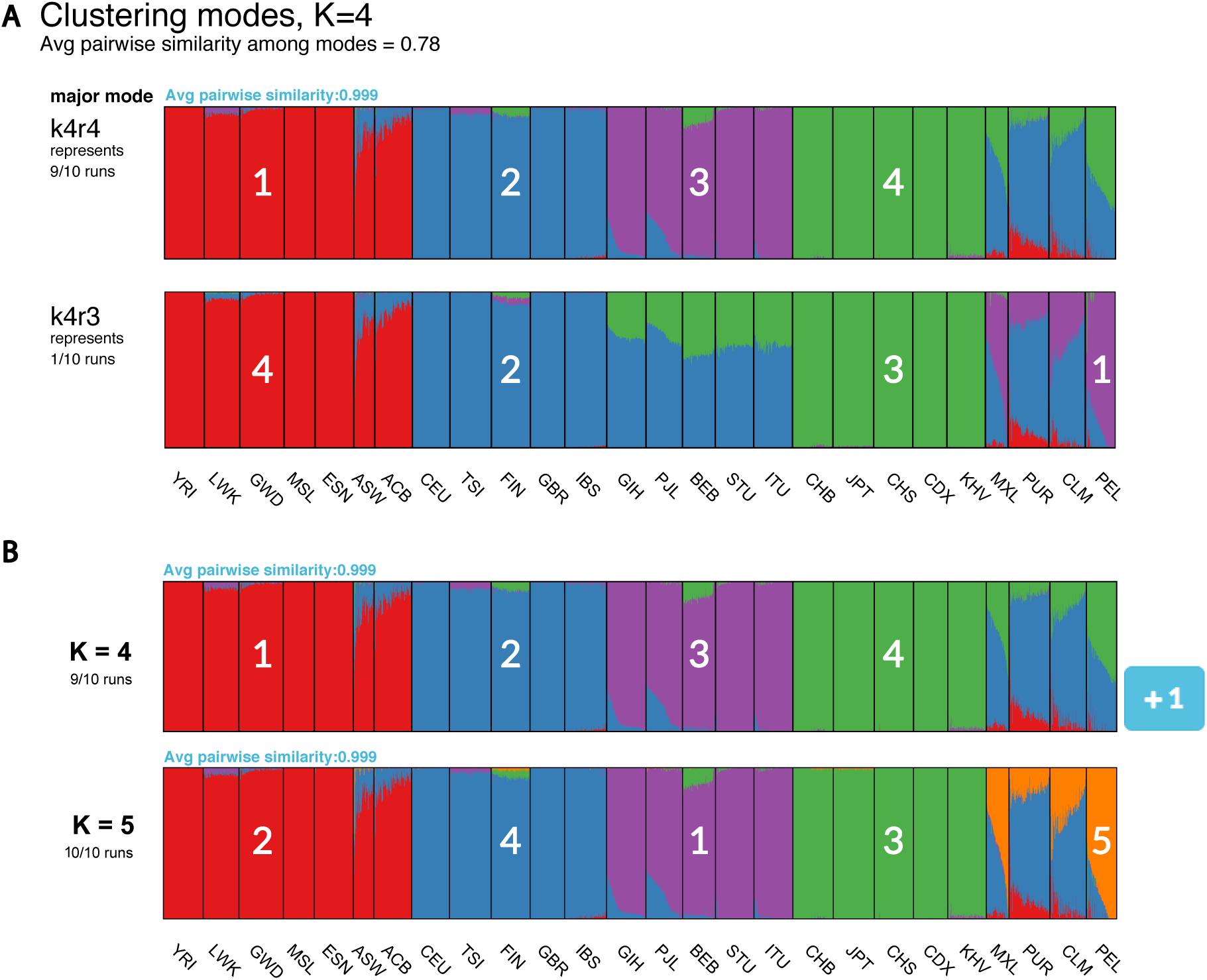
pong’s front end produces a D3.js visualization of maximum-weight alignments between runs, shown here for 20 *Q* matrices produced from clustering inference with ADMIXTURE (Alexander <u>et al.</u>, 2009) applied to 1000 Genomes data (phase 3; Consortium (2015)). Each individual’s genome-wide ancestry within a barplot is depicted by *K* stacked colored lines. The left-to-right order of individuals is the same in each barplot. The barplots here are annotated with numbers indicating which column of the underlying *Q* matrix is represented by a given cluster. **A:** Characterizing modes at *K* = 4 by displaying the representative run of the major mode (here, k4r4) and the representative run of each minor mode. Population codes are shown at the bottom. **B:** The maximum-weight alignment for the representative run for the major mode at *K* = 4 (k4r4, panel A) to that at *K* = 5. Membership in cluster 4 at *K* = 4 represents shared ancestry in East Asian and admixed American populations, and has been partitioned into Clusters 3 and 5 (representing East Asian and Native American ancestry, respectively) in the representative run of the major mode at *K* = 5.

We define an *alignment* of 𝓠 and 𝓡 as a bipartite perfect matching of their column vectors. `pong`’s first objective is to find the maximum-weight alignment for each pair of runs for a fixed value of *K* (Section 6.2). This information is used to identify modes within *K*, and we randomly choose a representative run (*Q* matrix) for each mode found in clustering inference. We call the mode containing the most runs within each value of *K* the *major mode* for that *K* value (Figure 1A; ties are decided uniformly at random). `pong`’s second objective is to find the maximum-weight alignment between the representative run of each major mode across values of *K* (Section 6.3; Figures 1B, S1). Identifying the maximum-weight alignment within and across *K* inherently solves the label switching problem without performing the computationally costly task of comparing whole-matrix permutations. Lastly, `pong` colors the visualization and highlights differences among modes based on these maximum-weight alignments.

### 6.2 Aligning runs for a fixed value of *K* and characterizing modes

In order to identify modes in clustering inference for a fixed value of *K* = *k*, `pong` first uses the Munkres algorithm (Munkres, 1957) to find the maximum-weight alignment between each pair of runs at *K* = *k* (Figure 2A). Next, for each value *k*, `pong` constructs another graph *G_k_* = ( {*Q*_*N* × *k*_ },*E*), where each edge connects a pair of runs, and the weight of a given edge is the average edge weight in the maximum-weight alignment for the pair of runs that edge connects. (The edge weight between a run and itself is 1.) The edge weight for a pair of runs in *G_k_* encodes the similarity of the runs, and we define *pairwise similarity* for a pair of runs as the average edge weight in the maximum-weight alignment across all clusters for that pair. We use the average edge weight to compute pairwise similarity instead of the sum of edge weights so that edges in *G_k_* are comparable across values of *K*.

**Figure 2:**
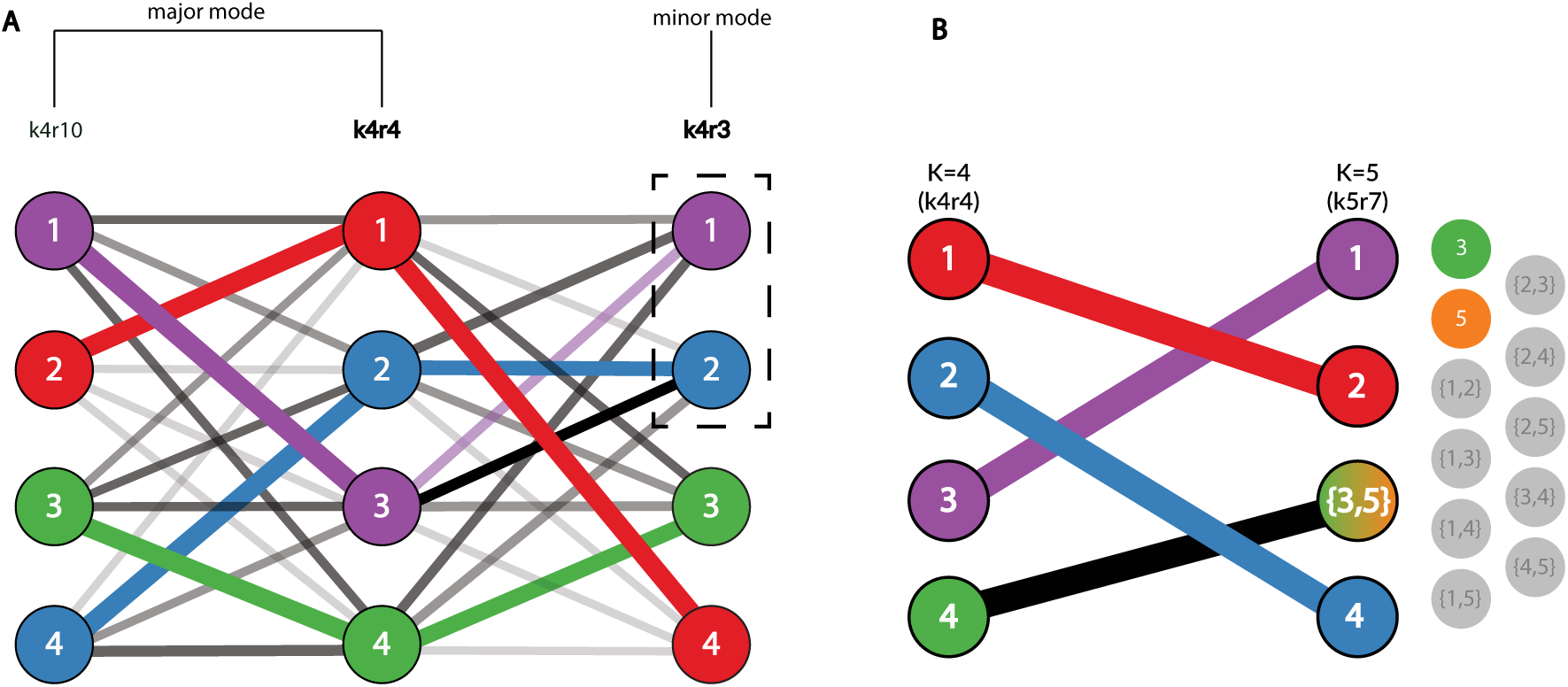
pong’s back-end model for the alignment of *Q* matrices, shown here from clustering inference with ADMIXTURE (Alexander <u>et al.</u>, 2009) applied to 1000 Genomes data (phase 3; Consortium (2015)). Panel labels correspond to panels in Figure 1, and numbers in graph vertices correspond to the clusters labeled in Figure 1. **A:** Characterizing modes from three runs of clustering inference at *K* = 4, the smallest *K* value with multiple modes for this dataset. Edge thickness corresponds to the value of `pong`’s default cluster similarity metric 𝓙 (derived from Jaccard’s index; see Supplementary Information), while edge opacity ranks connections for a cluster in run k4r4 to a cluster in run k4r3 (or in run k4r10). Note that both cluster 2 and 3 in k4r4 are most similar based on metric 𝓙 to cluster 2 in k4r3; in order to find the maximum-weight perfect matching between the runs, `pong` matches cluster 3 in k4r4 with cluster 1 in k4r3. Bold labels indicate representative runs for the two modes. Seven other runs (not displayed for ease of visualization) are grouped in the same mode as k4r4 and k4r10; these nine runs comprise the major mode at *K* = 4 (Figure 1A). k4r3 is the only run in the minor mo de (Figure 1A). **B:** Alignment of representative runs for the major modes at *K* = 4 to *K* = 5. 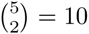 alignments are constructed between k4r4 and k5r7 (the representative run of the major mode at *K* = 5), constrained by the use of exactly one union node at *K* = 5. Of these 10 alignments, the alignment with maximum edge weight is shown and matches cluster 4 in k4r4 to the sum of clusters 3 and 5 in k5r7. The best matching for all other clusters are shown and informs the coloring of `pong`’s visualization (see Figure 1B).

If a pair of runs has pairwise similarity less than 0.97 (by default; this threshold can be varied), the edge connecting that pair of runs is not added to *G_k_*; this imposes a lower bound on the pairwise similarity between two runs in the same mode. `pong` defines modes as disjoint cliques in *G_k_*, thereby solving the multimodality problem. Our approach is informed by the fact that modes differ in only a subset of membership coefficients, eliminating the need for permuting whole matrices to align runs. Once cliques are identified, a run is chosen at random to be the representative run for each mode at *K* = *k*, which enables consistent visualization of clustering inference output within each value of *K*.

### 6.3 Aligning a *Q*_*N* × *K*_ matrix to a *Q*_*N* × (*K* * 1)_ matrix

Consider two *Q* matrices 𝓣_*N* × *k*_ and 𝓤_*N* × (*k* + * 1)_ where 𝓣 and 𝓤 represent the major modes at *K* = *k* and *K* = (*k* + *1), respectively. No perfect matching can be found between the clusters in 𝓣 and the clusters in 𝓤 because these matrices have different dimensions. In order to align these matrices, `pong` leverages the fact that column vectors of membership coefficients are partitioned as *K* increases and summed as *K* decreases (Figure 1B).

For the pair of clusters 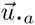 and 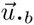 in 𝓤, we define the *union node* 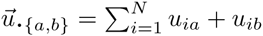. `pong` then constructs the matrix 𝓤(*a* ∪ *b*), which contains the clusters 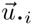 for *i* ≠ *a, b* and the union node 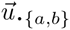. Therefore, the dimension of 𝓤(*a* ∪ *b*) is *N* × *K*, which is the same as the dimension of 𝓣 (Figure 2B). `pong` then finds the maximum-weight alignment between 𝓣 and 𝓤(*a* ∪ *b*) using the Munkres algorithm (Munkres, 1957). After finding the maximum-weight alignment for each pair of matrices 𝓣 and 𝓤(*i* ∪ *j*)(*i* ≠ *j*), the alignment that has the greatest average edge weight across all these 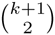 alignments is then used to solve the alignment-across-*K* problem. `pong` begins alignment across *K* between the representative runs of the major modes at *K* = 2 and *K* = 3 and proceeds through aligning *K* = *K_max_* – 1 and *K* = *K_max_*.

## 7 Implementation

`pong`’s back end is written in Python. While providing covariates is strongly advised so visualizations can be annotated with relevant metadata, `pong` only requires one tab-delimited file containing: (*i*) a user-provided identification code for each run (e.g., k4r4 in Figure 1A), (*ii*) the *K*-value for each run, and (*iii*) the relative path to each *Q* matrix. `pong` is executed with a one-line command in the terminal, which can contain a series of flags to customize certain algorithmic and visualization parameters. `pong`’s back end then generates results from its characterization of modes and alignment procedures that are printed to a series of output files.

After characterizing modes and aligning runs, `pong` initializes a local web server instance to host the visualization. `pong` is packaged with all its dependencies, such that it can be run without an Internet connection. The user is prompted to open a web browser and navigate to a specified port, and the user’s actions in the browser window lead to the exchange of data, such as *Q* matrices, via web sockets. These data are bound to and used to render the visualization.

pong’s front-end visualization is implemented in D3.js. `pong`’s main visualization displays the representative *Q* matrix for the major mode for each value of *K* as a Scalable Vector Graphic (SVG), where each individual’s genome-wide ancestry is depicted by *K* stacked colored lines. Each SVG is annotated with its value of *K*, the number of runs grouped into the major mode, and the average pairwise similarity across all pairs of runs in the major mode (Figure 1B).

For each value of *K*, a button is displayed to the right of the main visualization indicating the number of minor modes, if any exist (Figure 1B). Clicking on the button opens a pop-up dialog box consisting of barplots for the representative run of each mode within the *K* value, and each plot is annotated with the representative run’s user-provided identification code and the number of runs in each mode (Figure 1A). A dialog header reports the average pairwise similarity among pairs of representative runs for each mode, if there is more than one mode. Users can print or download any barplot in `pong`’s visualization in Portable Document Format (PDF) from the browser window.

What truly sets `pong`’s visualization apart from existing methods for the graphical display of population structure is a series of interactive features, which we now detail. In the browser’s main visualization, the user may click on any population — or set of populations, by holding SHIFT - to highlight the selected group’s genome-wide ancestry across values of K. When mousing over a population, the population’s average membership (as a percentage) in each cluster is displayed in a tooltip. Within each dialog box characterizing modes, selecting a checkbox on the top right allows the user to highlight differences between the major mode’s representative plot and each minor mode’s representative plot (Figure 3A). Clusters that do not differ beyond a threshold between a given major and minor mode are then shown as white in the minor mode, while the remaining clusters are shown at full opacity (Figure 3A; see also edge weights in Figure 2A).

**Figure 3:**
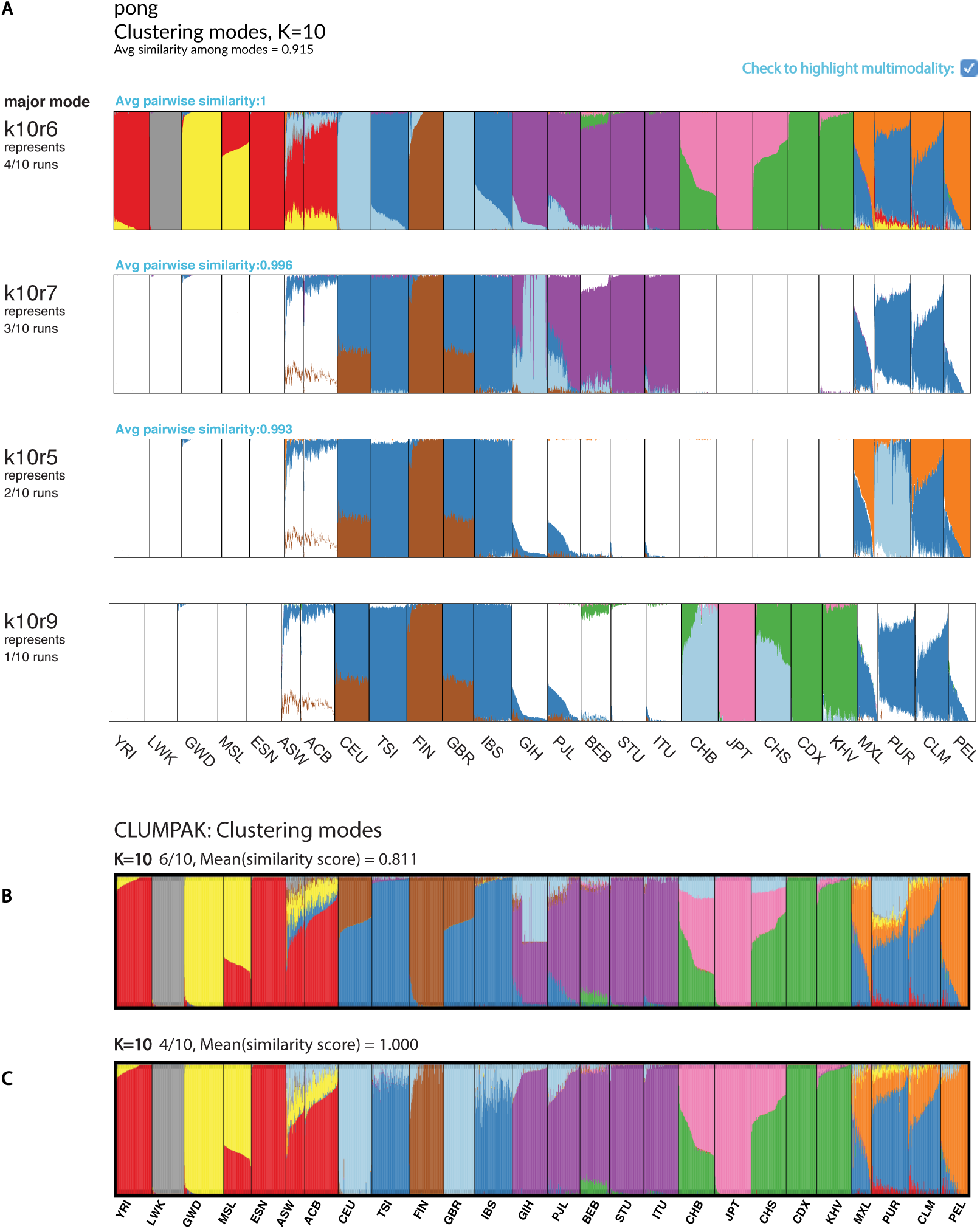
Visualizations of modes in population structure identified by `pong` and Clumpak at *K* =10 for clustering inference with ADMIXTURE (Alexander <u>et al.</u>, 2009) applied to 1000 Genomes data (phase 3; Consortium (2015)). The new cluster of membership coefficients first identified at *K* = 10 is denoted by light blue in each barplot. **A:** `pong`’s dialog box of modes at *K* = 10, with multimodality highlighted. **B:** Clumpak’s major mode at *K* = 10 averages over six runs of clustering inference output; the reported mean similarity score among these six runs is 0.811. South Asian (GIH), Han Chinese (CHB and CHS), and Puerto Rican (PUR) individuals all have ancestry depicted by the light blue cluster in this plot. The six runs averaged here are instead partitioned into three minor modes by `pong` in panel A. **C:** Clumpak’s minor mode at *K* =10 averages over four identical runs (mean similarity score is 1.000). This barplot contains the same information as the barplot of k4r10, representing `pong`’s major mode in panel A.

## 8 Results

We ran ADMIXTURE (Alexander <u>et al.</u>, 2009) on 225,705 unlinked genome-wide single-nucleotide variants from 2,426 unrelated individuals in the 1000 Genomes Project (phase 3, Consortium (2015); see Supplementary Information) to characterize population structure among globally distributed human populations. ADMIXTURE was run with *K* ranging from 2 to 10, and 10 runs were generated per value of *K*. Thus, a total of 90 *Q* matrices were produced; Figures 1 and 2 depict `pong`’s analysis of 20 of these runs. We also applied Clumpak (Kopelman <u>et al.</u>, 2015), the state-of-the-art method for automated post-processing and visualization of clustering inference output, to these 90 runs (partial results shown in Figures 3B,C; see also Supplementary Figure S2).

Clumpak automatically runs *CLUMPP* (Jakobsson and Rosenberg, 2007) for each value of *K* as part of its pipeline, and produces visualizations within and across values of *K* using Distruot (Rosenberg, 2004), displaying one barplot per mode. Figure 3B shows Clumpak’s reported major mode in the 1000 Genomes dataset at *K* = 10, which averages over six runs; all major modes reported by Clumpak can be viewed in Supplementary Figure S2. Using Clumpak’s web server (http://clumpak.tau.ac.il/) with its default settings (including using *CLUMPP*’s fastest algorithm, LargeKGreedy, for aligning *Q* matrices for a fixed value of *K*) took 58 minutes and 18 seconds for post-processing of these 90 runs. We could not apply other *CLUMPP* algorithms to the 1000 Genomes dataset using Clumpak’s web server due to the server’s restrictions against exhaustive running times (Kopelman <u>et al.</u>, 2015). We also installed Clumpak locally on Linux machines running Debian GNU/Linux 8 with 8 GB of RAM. Processing these 90 *Q* matrices took 74.275 hours using *CLUMPP*’s LargeKGreedy algorithm; using *CLUMPP*’s Greedy algorithm, which has increased accuracy over LargeKGreedy, Clumpak did not complete processing these *Q* matrices after four days. We also applied *CLUMPP*’s FullSearch algorithm, its most accurate algorithm, to the 10 *Q* matrices where *K* = 10; after 6.78 days, the job had still not completed.

Under its default settings, `pong` parsed input, characterized modes and aligned *Q* matrices within each value of *K*, and aligned *Q* matrices across *K* in 17.5 seconds on a Mid-2012 MacBook Pro with 8 GB RAM. After opening a web browser, `pong`’s interactive visualization loaded in an additional 3.2 seconds (Supplementary Figure S1 shows the main visualization).

In Figure 3A, `pong` identifies four modes at *K* = 10 in the 1000 Genomes dataset (phase 3). Light blue represents the cluster of membership coefficients first identified at *K* = 10 (see also Supplementary Figures S1, S2). In run k10r4 (representing 4 out of 10 runs), light blue represents British/Central European ancestry in the major mode (CEU and GBR). However, light blue represents South Asian ancestry (GIH) in 3 out of 10 runs (e.g., run k10r7), Puerto Rican ancestry (PUR) in 2 out of 10 runs (e.g., run k10r3), and Han Chinese ancestry in run k10r9. `pong`’s display of representative runs for each mode allows the user to observe and interpret multiple sets of membership coefficients inferred from the data at a given value of K.

In contrast, the minor mode Clumpak outputs (Figure 3C) is the same as `pong`’s major mode (Figure 3A), while Clumpak’s major mode reported at *K* = 10 (Figure 3B) averages over all minor modes identified by `pong`. The light blue in Clumpak’s reported major mode could be easily misinterpreted as shared ancestry among South Asian, East Asian, and Puerto Rican individuals, when in actuality these are distinct modes. We note that the highest-likelihood value of *K* for the 1000 Genomes data we analyzed is *K* = 8; at that value of *K*, we also see that Clumpak’s major mode suggests shared ancestry among individuals that are actually identified as having non-overlapping membership coefficients when individual runs are examined (Supplementary Figures S1, S2).

Figure 3A is the first visualization of some of the modes observed in population structure of 1000 Genomes phase 3 data, as Consortium (2015) ran ADMIXTURE exactly once per *K* value (see Extended Data Figure 5 and Supplementary Text of Consortium (2015)). Figure S3 shows `pong`’s visualization with consistent colors of all *Q* matrices released by Consortium (2015), *K* = 5 through 25; `pong` was able to process these *Q* matrices and render its visualization in 67.06 seconds. The modes identified in Figures 1, 3, and S1 differ substantially from the results reported by Consortium (2015). For example, in Figure 3A, `pong` depicts substructure in Puerto Rico and in China that is not observed in Extended Data Figure 5 by Consortium (2015). This could be due to different filters applied to the input SNP data (e.g., we removed relatives from data but did not filter based on minor allele frequency; see Supplementary Information), and we further note that these contrasting results indicate the need for efficient and accurate methods for processing and visualizing *Q* matrices.

## 9 Discussion

Here we introduce `pong`, a freely available user-friendly network-graphical method for post-processing output from clustering inference using population genetic data. We demonstrate that `pong` accurately aligns *Q* matrices orders of magnitude more quickly than do existing methods; it also provides a detailed characterization of modes among runs and produces a customizable, interactive D3.js visualization securely displayed using any modern web browser without requiring an internet connection. `pong`’s algorithm deviates from existing approaches by finding the maximum-weight perfect matching between column vectors of membership coefficients for pairs of *Q* matrices, and leverages the Hungarian algorithm to efficiently solve this series of optimization problems (Kuhn, 1955, 1956; Munkres, 1957).

Interpreting the results from multiple runs of clustering inference is a difficult process. Investigators often choose a single *Q* matrix at each value of *K* to display or discuss, overlooking complex signals present in their data because the process of producing the necessary visualizations is too time-consuming. `pong`’s speed allows the investigator to focus instead on conducting more runs of clustering inference in order to fully interpret the clustering in her sample of interest. Currently, many population-genetic studies only carry out one run of clustering inference per value of *K* (Moreno-Estrada <u>et al.</u>, 2013; Consortium, 2015; Homburger <u>et al.</u>, 2015; Mathieson <u>et al.</u>, 2015; Lorenzi <u>et al.</u>, 2016; Hallast <u>et al.</u>, 2016; Jeffares <u>et al.</u>, 2015), particularly when using ADMIXTURE’s maximum-likelihood approach (Alexander <u>et al.</u>, 2009) to the inferential framework implemented in Structure (Pritchard <u>et al.</u>, 2000). The likelihood landscape of the input genotype data is complex, and can hold different local maxima for a given value of *K* (see Verdu <u>et al.</u> (2014)). Combining `pong`’s rapid algorithm and detailed, interactive visualization with posterior probabilities for *K* reported by clustering inference methods will allow investigators to accurately interpret results from clustering inference, thereby advancing our knowledge of the genetic structure of natural populations for a wide range of organisms. We further plan to extend `pong` to visualize results from other applications of mixed-membership models and to leverage the dynamic nature of bound data to increase the information provided by `pong`’s visualization.

## Acknowledgements

We thank the Ramachandran Lab for helpful discussions, and three anonymous reviewers for their comments on an earlier version of this manuscript. Data sets for testing early versions of `pong` were provided by Elizabeth Atkinson and Brenna Henn, Charleston Chiang and John Novembre, Caitlin Uren, and Paul Verdu; the Henn and Novembre labs, Chris Gignoux, and Catherine Luria tested beta versions of `pong`. We also thank Mark Howison for helpful discussions regarding python packaging. Multiple members of the Raphael Lab at Brown University helped improve `pong`, especially Max Leiserson and Hsin-Ta Wu, who advised on D3.js, and Mohammed El-Kebir, whose suggestions increased the efficiency of the back end and improved the manuscript.

## Cluster similarity metrics

We implemented and tested several metrics for cluster similarity. The default metric used by `pong`, 𝓙 (Equation 1), is derived from the Jaccard index used in set comparison. For a given pair of clusters 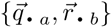, let *N*\* be the set of indices for which at least one of 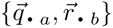 has a nonzero entry; that is, *N*\* = {*i* ∈ {1,…, *N* }: *q_ia_* * *r_ib_* > 0 }. Then,

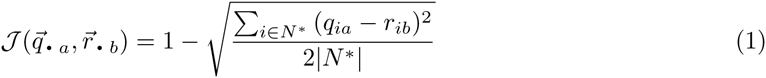

𝓙 is designed to emphasize overlap in membership coefficients while ignoring overlap in nonmembership (i.e., individuals with membership coefficients of 0 in the clusters under comparison). Although we recommend using 𝓙, `pong` implements other similarity metrics: *G*′ (as used in *CLUMPP* Jakobsson and Rosenberg (2007)), the average sum of squared differences between *q.* 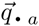 and 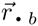 (subtracted from 1), and average Manhattan distance (subtracted from 1). `pong`’s implementation is designed such that users familiar with Python and NumPy can add their own similarity metrics to the source code if desired.

## Processing of 1000 Genomes Data

Variant calls for 1,019,196 genome-wide single-nucleotide variants (SNVs) in 2,504 individuals were extracted from the 1000 Genomes Project Phase 3 data repository ftp://ftp.1000genomes.ebi.ac.uk/vol1/ftp/release/20130502/ (release date: Nov 6, 2014), using the command-line tool `tabix` (Li, 2011).

A total of 78 individuals were excluded from analysis based on relatedness: one individual from each pair of first- and second-degree relatives was removed, leaving a total of 2,426 individuals. Next, SNVs were pruned for linkage disequilibrium using the `-indep-pairwise` flag in PLINK (Purcell <u>et al.</u>, 2007). We removed every SNV with *r*^2^ > 0.1 with any other SNV within a 50-SNV sliding window (PLINK command-line parameters for `-indep-pairwise`: 50 10 0.01), leaving a total of 225,705 SNVs for analysis.

ADMIXTURE (Alexander <u>et al.</u>, 2009) was applied 10 times per value of *K* to these data, with *K* taking on values in the closed interval [2,10]. The value of *K* that minimized cross-validation error was *K* = 8.

---

**Figure S1:**
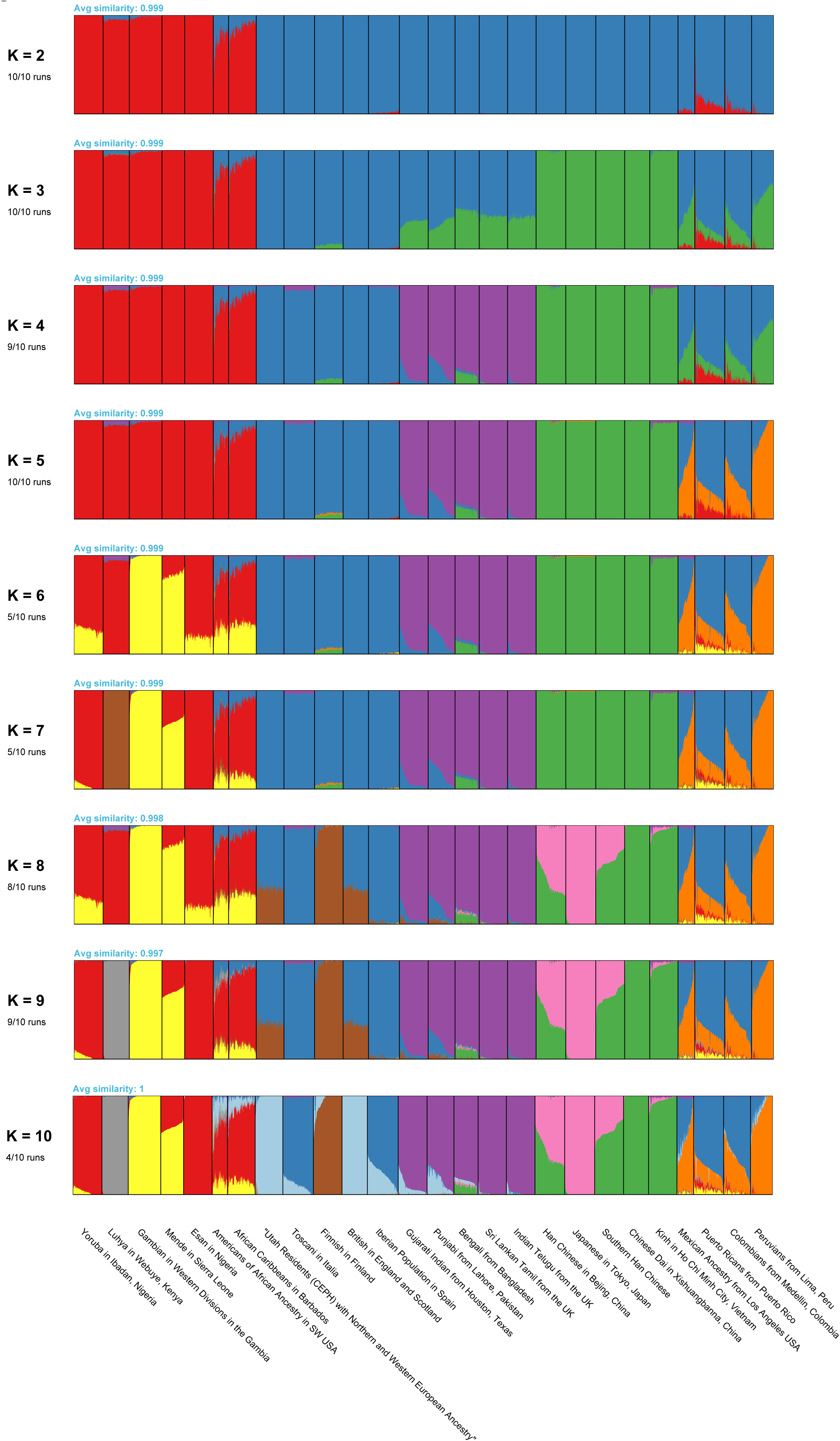
`pong`’s main visualization of major modes in population structure in the 1000 Genomes (phase 3) with detailed population labels, *K* = 2 through *K* = 10.

**Figure S2:**
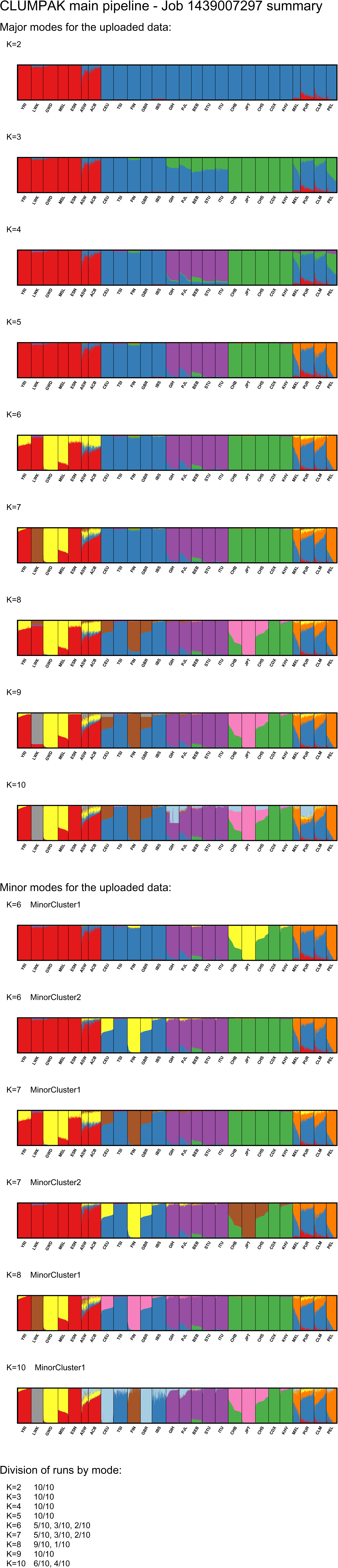
Clumpak’s (Kopelman <u>et al.</u>, 2015) visualization of modes in population structure in the 1000 Genomes (phase 3), *K* = 2 through *K* = 10.

**Figure S3:**
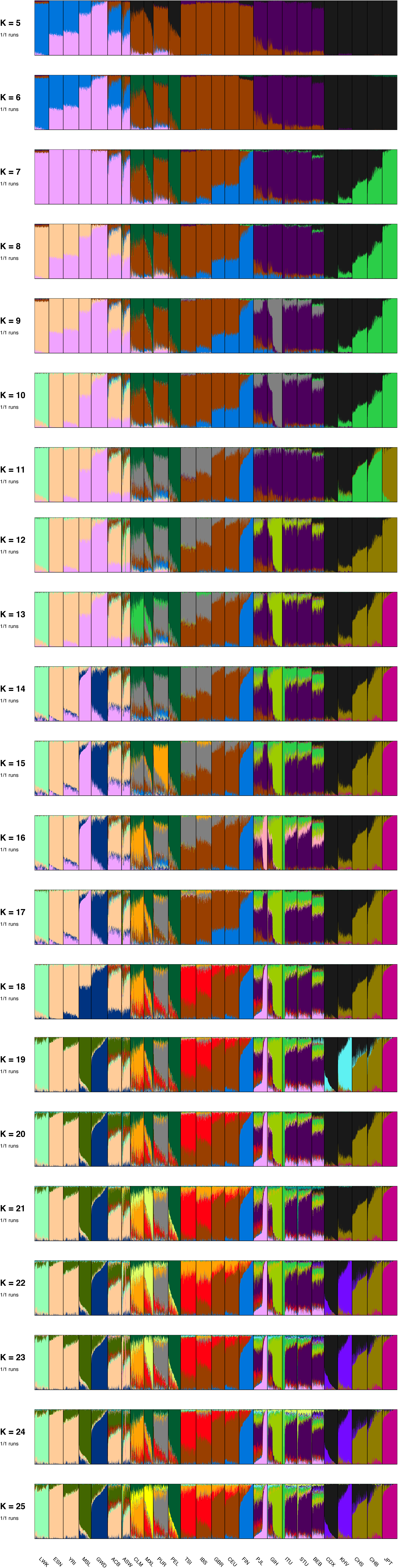
`pong`’s visualization of population structure in the 1000 Genomes (phase 3), based on *Q* matrices from Consortium (2015), *K* = 5 through *K* = 25.

## References

Alexander, D. H., Novembre, J., and Lange, K. (2009). Fast model-based estimation of ancestry in unrelated individuals. Genome Research, 19(9), 1655–1664.

Blei, D. M., Ng, A. Y., and Jordan, M. I. (2003). Latent Dirichlet Allocation. Journal of Machine Learning Research, 3(4-5), 993–1022.

Bryc, K., Auton, A., Nelson, M. R., Oksenberg, J. R., Hauser, S. L., Williams, S., Froment, A., Bodo, J.-M., Wambebe, C., Tishkoff, S. a., and Bustamante, C. D. (2010). Genome-wide patterns of population structure and admixture in West Africans and African Americans. Proceedings of the National Academy of Sciences of the United States of America, 107(2), 786–91.

Consortium, T.. G. P. (2015). A global reference for human genetic variation. Nature, 526(7571), 68–74.

Falush, D., Stephens, M., and Pritchard, J. K. (2003). Inference of population structure using multilocus genotype data: linked loci and correlated allele frequencies. Genetics, 164(4), 1567–87.

Galanter, J. M., Fernandez-Lopez, J. C., Gignoux, C. R., Barnholtz-Sloan, J., Fernandez-Rozadilla, C., Via, M., Hidalgo-Miranda, A., Contreras, A. V., Figueroa, L. U., Raska, P., Jimenez-Sanchez, G., Zolezzi, I. S., Torres, M., Ponte, C. R., Ruiz, Y., Salas, A., Nguyen, E., Eng, C., Borjas, L., Zabala, W., Barreto, G., González, F. R., Ibarra, A., Taboada, P., Porras, L., Moreno, F., Bigham, A., Gutierrez, G., Brutsaert, T., León-Velarde, F., Moore, L. G., Vargas, E., Cruz, M., Escobedo, J., Rodriguez-Santana, J., Rodriguez-Cintrón, W., Chapela, R., Ford, J. G., Bustamante, C., Seminara, D., Shriver, M., Ziv, E., Burchard, E. G., Haile, R., Parra, E., and Carracedo, A. (2012). Development of a panel of genome-wide ancestry informative markers to study admixture throughout the americas. PLoS Genetics, 8(3), e1002554.

Glover, K. a., Quintela, M., Wennevik, V., Besnier, F., Sø rvik, A. G. E., and Skaala, O. y. (2012). Three decades of farmed escapees in the wild: A spatio-temporal analysis of atlantic salmon population genetic structure throughout norway. PLoS ONE, 7(8), e43129.

Hallast, P., Delser, P. M., Batini, C., Zadik, D., Schempp, W., Tyler-smith, C., and Jobling, M. A. (2016). Great-ape Y-Chromosome and mitochondrial DNA phylogenies reflect sub-species structure and patterns of mating and dispersal Corresponding author:. Genome research, 44(0), 1–13.

Homburger, J. R., Moreno-Estrada, A., Gignoux, C. R., Nelson, D., Sanchez, E., Ortiz-Tello, P., Pons-Estel, B. A., Acevedo-Vasquez, E., Miranda, P., Langefeld, C. D., Gravel, S., Alarcón-Riquelme, M. E., and Bustamante, C. D. (2015). Genomic Insights into the Ancestry and Demographic History of South America. PLoS Genetics, 11(12), 1–26.

Hubisz, M. J., Falush, D., Stephens, M., and Pritchard, J. K. (2009). Inferring weak population structure with the assistance of sample group information. Molecular Ecology Resources, 9(5), 1322–1332.

Huelsenbeck, J. P., Andolfatto, P., and Huelsenbeck, E. T. (2011). Structurama: bayesian inference of population structure. Evolutionary Bioinformatics, 7, 55–9.

Jakobsson, M. and Rosenberg, N. a. (2007). CLUMPP: a cluster matching and permutation program for dealing with label switching and multimodality in analysis of population structure. Bioinformatics (Oxford, England), 23(14), 1801–6.

Jasra, A., Holmes, C., and Stephens, D. (2005). Markov Chain Monte Carlo methods and the label switching problem in Bayesian Mixture Modeling. Statistical Science, 20(1), 50–67.

Jeffares, D. C., Rallis, C., Rieux, A., Speed, D., PÅŹevorovský, M., Mourier, T., Marsellach, F. X., Iqbal, Z., Lau, W., Cheng, T. M. K., Pracana, R., Mülleder, M., Lawson, J. L. D., Chessel, A., Bala, S., Hellenthal, G., O’Fallon, B., Keane, T., Simpson, J. T., Bischof, L., Tomiczek, B., Bitton, D. A., Sideri, T., Codlin, S., Hellberg, J. E. E. U., van Trigt, L., Jeffery, L., Li, J.-J., Atkinson, S., Thodberg, M., Febrer, M., McLay, K., Drou, N., Brown, W., Hayles, J., Carazo Salas, R. E., Ralser, M., Maniatis, N., Balding, D. J., Balloux, F., Durbin, R., and Bähler, J. (2015). The genomic and phenotypic diversity of Schizosaccharomyces pombe. Nature genetics, 47(3), 235–41.

Kopelman, N. M., Mayzel, J., Jakobsson, M., Rosenberg, N. A., and Mayrose, I. (2015). C LUMPAK: a program for identifying clustering modes and packaging population structure inferences across K. Molecular Ecology Resources, pages doi: 10.1111/1755-0998.12387.

Kuhn, H. W. (1955). The Hungarian Method for the assignment problem. Naval Research Logistics Quarterly, 2, 83–97.

Kuhn, H. W. (1956). Variants of the Hungarian method for assignment problems. Naval Research Logistics Quarterly, 3, 253–258.

Lorenzi, H., Khan, A., Behnke, M. S., Namasivayam, S., Swapna, L. S., Hadjithomas, M., Karamycheva, S., Pinney, D., Brunk, B. P., Ajioka, J. W., Ajzenberg, D., Boothroyd, J. C., Boyle, J. P., Dardé, M. L., Diaz-Miranda, M. a., Dubey, J. P., Fritz, H. M., Gennari, S. M., Gregory, B. D., Kim, K., Saeij, J. P. J., Su, C., White, M. W., Zhu, X.-Q., Howe, D. K., Rosenthal, B. M., Grigg, M. E., Parkinson, J., Liu, L., Kissinger, J. C., Roos, D. S., and David Sibley, L. (2016). Local admixture of amplified and diversified secreted pathogenesis determinants shapes mosaic Toxoplasma gondii genomes. Nature Communications, 7, 10147.

Manber, U. (1989). Introduction to Algorithms: A Creative Approach. Addison-Wesley.

Mathieson, I., Lazaridis, I., Rohland, N., Mallick, S., Patterson, N., Roodenberg, S. A., Harney, E., Stewardson, K., Fernandes, D., Novak, M., Sirak, K., Gamba, C., Jones, E. R., Llamas, B., Dryomov, S., Pickrell, J., Arsuaga, J. L., de Castro, J. M. B., Carbonell, E., Gerritsen, F., Khokhlov, A., Kuznetsov, P., Lozano, M., Meller, H., Mochalov, O., Moiseyev, V., Guerra, M. A. R., Roodenberg, J., Vergès, J. M., Krause, J., Cooper, A., Alt, K. W., Brown, D., Anthony, D., Lalueza-Fox, C., Haak, W., Pinhasi, R., and Reich, D. (2015). Genome-wide patterns of selection in 230 ancient Eurasians. Nature, 528(7583), 499–503.

Moore, A. J., Moore, W. L., and Baldwin, B. G. (2014). Genetic and ecotypic differentiation in a californian plant polyploid complex (Grindelia, Asteraceae). PLoS ONE, 9(4), e95656.

Moreno-Estrada, A., Gravel, S., Zakharia, F., McCauley, J. L., Byrnes, J. K., Gignoux, C. R., Ortiz-Tello, P. a., Martínez, R. J., Hedges, D. J., Morris, R. W., Eng, C., Sandoval, K., Acevedo-Acevedo, S., Norman, P. J., Layrisse, Z., Parham, P., Martínez-Cruzado, J. C., Burchard, E. G., Cuccaro, M. L., Martin, E. R., and Bustamante, C. D. (2013). Reconstructing the population genetic history of the Caribbean. PLoS Genetics, 9(11), e1003925.

Moreno-Estrada, A., Gignoux, C. R., Fernández-López, J. C., Zakharia, F., Sikora, M., Contreras, A. V., Acuña Alonzo, V., Sandoval, K., Eng, C., Romero-Hidalgo, S., Ortiz-Tello, P., Robles, V., Kenny, E. E., Nuño Arana, I., Barquera-Lozano, R., Macín-Pérez, G., Granados-Arriola, J., Huntsman, S., Galanter, J. M., Via, M., Ford, J. G., Chapela, R., Rodriguez-Cintron, W., Rodríguez-Santana, J. R., Romieu, I., Sienra-Monge, J. J., del Rio Navarro, B., London, S. J., Ruiz-Linares, A., Garcia-Herrera, R., Estrada, K., Hidalgo-Miranda, A., Jimenez-Sanchez, G., Carnevale, A., Soberón, X., Canizales-Quinteros, S., Rangel-Villalobos, H., Silva-Zolezzi, I., Burchard, E. G., and Bustamante, C. D. (2014). The genetics of Mexico recapitulates Native American substructure and affects biomedical traits. Science (New York, N.Y.), 344(6189), 1280–5.

Munkres, J. (1957). Algorithms for the Assignment and Transportation Problems. Journal of the Society of Industrial and Applied Mathematics, 5(1), 32–38.

Novembre, J. (2014). Variations on a common STRUCTURE: new algorithms for a valuable model. Genetics, 197(3), 809–811.

Patterson, N., Price, A. L., and Reich, D. (2006). Population structure and eigenanalysis. PLoS Genetics, 2(12), e190.

Price, A. L., Patterson, N. J., Plenge, R. M., Weinblatt, M. E., Shadick, N. a., and Reich, D. (2006). Principal components analysis corrects for stratification in genome-wide association studies. Nature Genetics, 38(8), 904–909.

Pritchard, J. K., Stephens, M., and Donnelly, P. (2000). Inference of population structure using multilocus genotype data. Genetics, 155(2), 945–959.

Raj, A., Stephens, M., and Pritchard, J. K. (2014). fastSTRUCTURE: Variational Inference of Population Structure in Large SNP Data Sets. Genetics, 197(2), 573–589.

Rosenberg, N. a. (2004). DISTRUCT: A program for the graphical display of population structure. Molecular Ecology Notes, 4(1), 137–138.

Stephens, M. (2000). Dealing with label switching in mixture models. J. R. Statist. Soc. Series B, 62(4), 795–809.

Verdu, P., Pemberton, T. J., Laurent, R., Kemp, B. M., Gonzalez-Oliver, A., Gorodezky, C., Hughes, C. E., Shattuck, M. R., Petzelt, B., Mitchell, J., Harry, H., William, T., Worl, R., Cybulski, J. S., Rosenberg, N. a., and Malhi, R. S. (2014). Patterns of Admixture and Population Structure in Native Populations of Northwest North America. PLoS genetics, 10(8), e1004530.

## References

Li, H. (2011). Tabix: fast retrieval of sequence features from generic TAB-delimited files. Bioinformatics, 27, 718–9.

Purcell, S., Neale, B., Todd-Brown, K., Thomas, L., Ferreira, M. A., Bender, D., Maller, J., Sklar, P., De Bakker, P. I., Daly, M. J., and Sham, P. C. (2007). PLINK: a tool set for whole-genome association and population-based linkage analyses. The American Journal of Human Genetics, 81(3), 559–575.

